# Spatiotemporal dynamics of NF-κB/Dorsal inhibitor IκBα/Cactus in *Drosophila* blastoderm embryos

**DOI:** 10.1101/2024.02.23.581825

**Authors:** Allison E. Schloop, Sharva Hiremath, Razeen Shaikh, Cranos M. Williams, Gregory T. Reeves

## Abstract

The NF-κB signaling pathway is a key regulatory network in mammals that controls many cellular processes, including immunity and inflammation. Of particular note is the relationship between NF-κB and its inhibitor IκBα, which sequesters NF-κB to the cytoplasm of cells until needed. It is also known that IκBα can enter nuclei, disrupt NF-κB binding to DNA, and shuttle it out to again sequester NF-κB in the cytoplasm. In *Drosophila melanogaster*, a homologous system between the proteins Dorsal (homologous to NF-κB) and Cactus (homologous to IκBα) is important in embryo development, specifically in establishment of the Dorsal nuclear concentration gradient. Previous work suggests Cactus also enters the nucleus; mathematical models of the Dorsal gradient fail to accurately predict the normal range of the gradient without nuclear Cactus. However, direct, *in vivo* visualization of Cactus spatiotemporal dynamics, including its localization to the nuclei, has been difficult to gather. Previously, imaging Cactus in live embryos was complicated by rapid protein turnover, preventing fluorescent protein fusions from fully maturing. To address this, we used the CRISPR/Cas9 system to tag Cactus with the recently developed “LlamaTag” (LT), a genetically encodable nanobody from llamas that dynamically binds to GFP *in vivo*. We then employed standard confocal imaging, as well as advanced optical techniques such as raster image correlation spectroscopy (RICS) and fluorescent recovery after photobleaching (FRAP) to investigate the spatiotemporal distribution of Cactus-LlamaTag in *Drosophila* embryos at the blastoderm stage. Our results demonstrate that Cactus can be found in the nuclei of early embryos, consistent with its role as a transcription factor regulator. Moreover, by using the data from FRAP and RICS, we were able to estimate biophysical parameters of Cactus dynamics *in vivo*, including its nuclear transport rate constants and fraction bound to GFP. These data were further used to constrain a mathematical model that allowed us to infer experimentally inaccessible biophysical parameters, such as the concentration of Cact protein and the dissociation constant of LT and GFP. Our study provides new insights into the regulation of the NF-κB pathway in early *Drosophila* embryos and highlights the power of advanced optical techniques for investigating complex biological dynamics.

## Introduction

Metazoan development is dependent upon proper regulation of gene expression, which often relies on long-range signals known as morphogens. These morphogens form concentration gradients that dictate spatially-dependent expression of genes responsible for proper body patterning (Driever & Nüsslein-Volhard, 1988a, 1988b; Roth et al., 1989; Rushlow et al., 1989; Steward, 1989; Struhl et al., 1989). In *Drosophila*, the transcription factor Dorsal (Dl), a homolog of vertebrate NF-κB, acts as a morphogen to pattern the dorsal-ventral (DV) axis in the early (1-3 h old) embryo (Roth et al., 1989; Rushlow et al., 1989; Schloop et al., 2020; Steward, 1989). Cactus (Cact), a homolog of vertebrate IκBα, is initially bound to Dl, sequestering it to the cytoplasm. Toll signaling on the ventral side of the embryo degrades Cact and allows Dl to enter the nucleus to regulate target gene expression (Hashimoto et al., 1991; Roth et al., 1989; Rushlow et al., 1989; Schneider et al., 1991; Stein et al., 1991; Steward et al., 1988). Ultimately, Dl activity specifies multiple gene expression domains along the DV axis to initialize the embryo fatemap (Chopra & Levine, 2009; Reeves et al., 2012; Reeves & Stathopoulos, 2009; Stathopoulos & Levine, 2004).

It is widely accepted that Cact acts as an inhibitor of Dl by binding to it and keeping it sequestered in the cytoplasm (Roth et al., 1991). This role was established during early research into dorsoventral polarity of *Drosophila* embryos, which also established its sequence similarity to IκB (Geisler et al., 1992). Further work determined that establishment of dorsoventral polarity required a gradient of Cact degradation, as evidenced by antibody staining of Cact in the cytoplasm of the syncytial blastoderm (Bergmann et al., 1996; Reach et al., 1996). Taken together, this provided definitive evidence for the role of Cact in the cytoplasm. However, recent evidence suggests that, beyond cytoplasmic sequestration, Cact may also play a role in regulation of the Dorsal gradient from within the nucleus. Recent modeling work suggests that Cact must be present in the nucleus to explain both the dynamics of the Dl gradient as observed in live imaging (O’Connell & Reeves, 2015; Reeves et al., 2012), as well as the extent and robustness of Dl-dependent gene expression (Al Asafen et al., 2020). The presence of Cact in the nucleus was also inferred from a study measuring the mobility of Dl-GFP in live embryos (Al Asafen et al., 2018). Furthermore, in vertebrates, the Cact homolog IκBα also plays a role within the nucleus by entering the nucleus and disrupting NF-κB binding with DNA, which forces the NF-κB/IκBα complex to be exported back to the cytoplasm via a nuclear export signal (Arenzana-Seisdedos et al., 1995, 1997). Given the similarity between Cact and IκBα, it is reasonable to ask whether Cact may enter the nucleus.

Obtaining direct experimental evidence of nuclear Cact through established imaging techniques has proved to be a challenge. While two studies in the mid-1990s successfully used in situ immunostaining to image a weak Cact gradient, the results were not quantifiable and have not been replicated since (Bergmann et al., 1996; Reach et al., 1996). Moreover, immunostaining in fixed embryos provides only a snapshot of a highly dynamic system (DeLotto et al., 2007; Liberman et al., 2009; Reeves et al., 2012). On the other hand, live imaging typically requires tagging a protein of interest with a fluorescent protein. Development of the fly embryo happens over a matter of hours, with some crucial proteins performing their job in a matter of minutes (Edgar et al., 1987), and the turnover of developmental proteins during the *Drosophila* blastoderm stage is often too rapid to allow maturation of the fluorescent protein tag. In particular, Cact has a high turnover rate (Kidd, 1992). To circumvent this issue, here we are using a recently developed method referred to as LlamaTags to visualize Cact in a live setting (Bothma et al., 2018; Kirchhofer et al., 2010).

LlamaTags (LT) are nanobodies, small single-domain antibodies, that are derived from llamas. These LTs are genetically encodable and can be raised against particular fluorescent molecules and be fused to a protein of interest instead of using a fluorescent protein fusion(Bothma et al., 2018; Kirchhofer et al., 2010). In a recent *Drosophila* blastoderm study, the corresponding fluorescent protein (GFP or mCherry) was provided maternally, which resulted in the presence of mature fluorescent protein during the stage of interest (Bothma et al., 2018). This method has been explored with several transcription factors important for development, including Twist and Hunchback (Hb), with results of equivalent quality to previously used fluorescent protein fusions. For example, in embryos co-expressing GFP maternally and a fusion of Hb with the LT that binds GFP (Hb-LT) zygotically, GFP fluorescence recapitulated the known Hb expression pattern, including localization to the nuclei (Bothma et al., 2018). However, given that maternally provided GFP is expressed uniformly in the embryo, this method is subject to background GFP fluorescence that is not bound to the LT-fusion of interest. Therefore, this technique works with proteins that have specific subcellular localization patterns. For example, in LT fusions with transcription factors, which naturally localize to the nuclei, GFP fluorescence increases in the nuclei upon binding to the LT fusion (Bothma et al., 2018).

In this paper, we used an LT that binds GFP to measure the spatiotemporal localization and dynamics of Cact (Bothma et al., 2018; Kirchhofer et al., 2010). We used CRISPR to fuse the LT in-frame at the C-terminus of native Cact. In the homozygous state, the fly line expressing the Cact-LT CRISPR construct was healthy, viable, fertile, and did not perturb the Dl gradient, even in the presence of maternally-provided GFP. We visualized the nucleocytoplasmic localization of Cact in early (1-3 h old) embryos and measured its distribution along the DV axis. We found that, while Cact does not exhibit a DV gradient, it displays temporal dynamics similar to Dl (Reeves et al., 2012). We also used fluorescence recovery after photobleaching (FRAP) and raster image correlation spectroscopy (RICS) to determine that Cact is present in the nuclei. Using a steady state model of Cact-LT/GFP interactions, we estimated the concentration of Cact in the embryo and the affinity of GFP for the LT. Our results have implications for proper interpretations of measurements of the Dl gradient.

## Methods

### Fly lines

The Cactus-LlamaTag (Cact-LT) fly line was developed following the CRISPR protocol described by Gratz et al. 2015 (Gratz et al., 2015). Briefly, the sequence of GFP Llama-Tag was inserted into the pHD-DsRed plasmid such that it was a part of the right homology arm for homology directed repair (see Table S1 for gBlock sequence). gRNAs were designed to cut immediately before the native Cactus stop codon to insert the Llama-Tag, and in the closest intron for insertion of a red fluorescent eye marker for successful insertion selection (see Table S2 for primers). All plasmids were assembled via standard cloning techniques. Plasmids used in this protocol were gifted from Kate O’Connor-Giles (Addgene plasmids #45946 and #51434)(Gratz et al., 2013, 2014). The sequence of the GFP Llama-Tag was provided by Hernan Garcia (Bothma et al., 2018; Kirchhofer et al., 2010). Transgenic flies were generated by GenetiVision Corporation (Stafford, TX). The maternal eGFP fly line was also generously provided by Hernan Garcia (Kim et al., 2022).

The baseline Cact-LT line was Cact-LT/CyO; H2ARFP/TM3, crossed with +/+;eGFP/eGFP, to arrive at Cact-LT/+; H2ARFP/eGFP (1×1x). Experiments requiring 2 copies of eGFP used Cact-LT/CyO; eGFP/TM3 as a baseline and were crossed with H2ARFP/CyO; eGFP/TM3, for a final genotype of Cact-LT/H2ARFP; eGFP/eGFP (2x GFP). Experiments requiring 2 copies of Cact-LT used Cact-LT/CyO; H2ARFP/TM3 as a baseline and were crossed with Cact-LT/CyO; eGFP/TM3, with a resulting genotype of Cact-LT/Cact-LT; H2ARFP/eGFP (2x Cact-LT). Experiments requiring free GFP as a control used the fly line H2ARFP/CyO; eGFP/TM3 (control).

### Sample preparation for live imaging

Female offspring with appropriate genotype were selected and put in a collection cage with a group of males. These flies were allowed to lay eggs on grape juice plates for one hour at room temperature before being incubated at 25°C for 30 minutes to an hour for appropriate staging. Embryos were washed off the plate with deionized water into embryo collection baskets, bleached for 35-40 seconds to remove chorions, and then rinsed in deionized water. For whole-mount imaging, embryos were adhered to microscope slides using either heptane glue or low-melt agarose, immersed in water, and then covered with a cover slip. Halocarbon oil 27 was also used for whole-mount imaging, however, embryos were immersed solely in halocarbon oil before being covered with a cover slip. For end-on mounting, embryos were embedded in a drop of low melt agarose and oriented such that their anterior-posterior axis was perpendicular to a glass bottom petri dish (MatTek, No. 1.5 Coverslip, 20mm diameter well, P35G-1.5-20-C). After the agarose set, deionized water was added to the dish to prevent drying of the agarose.

### Imaging of Cross Sections of Live Embryos

To image optical cross sections of live embryos, we employed end-on mounting as described above. To capture a representative slice of the embryo, images were taken 150uM into the embryo from the end facing the lens. Images were taken every 2 minutes at a scan speed of 5 with 4x averaging, starting at nuclear cycle (nc) 10 or 11 and continuing until the embryo gastrulated. Laser power was kept below 5% to prevent photobleaching.

### Analysis of Cross Sections of Live Embryos

After optical cross sections of live, end-on mounted embryos were collected, the cross sections were analyzed according to the following methods. First, the nuclei and cytoplasmic compartments were segmented as described previously (Al Asafen et al., 2020; Trisnadi et al., 2013). To ensure fluorescent intensities of the cytoplasmic compartments were not influenced by light scattering due to the depth of imaging, the cytoplasmic compartments were constrained to lie between the apical side of the nuclear layer and the embryo periphery. Individual nuclei were assigned the DV coordinate of their centroid. Cytoplasmic compartments were separated by proximity to nuclei and were assigned the DV coordinate of the closest nucleus. The ventral midline, defined as the location of the ventral furrow at gastrulation, was assigned the DV coordinate of zero. The DV coordinate axis was defined as ranging from -1 to +1, with zero signifying the ventral midline and both -1 and +1 signifying the dorsal midline.

Average intensity values were extracted for each segmented nucleus and cytoplasmic compartment. These intensity values were filtered by hard thresholds to identify possible outliers, which were further investigated for segmentation errors. After the elimination of true outliers, possible variations in nuclear or cytoplasmic intensities across the DV axis were examined by a t-test on the slope of the best-fit line for each frame.

After extracting the intensities from the live embryos, an averaged plot for the behavior of Cactus across the cross section was generated. This plot was generated by first aligning the curves from the different embryos along their peaks and then using scaling to ensure that they have the same amplitudes. Finally, the mean of the intensities of nuclei and cytoplasm was calculated to generate the averaged embryo plot. For more information, see Supplementary Information.

### Imaging of Live Embryos using Fluorescent Recovery After Photobleaching (FRAP)

FRAP was performed on embryos of 3 fly lines: 1×1x, 2x GFP, and the control. Each line was prepared as noted for whole body imaging. A Zeiss LSM 900 confocal microscope was used to bleach a specific nucleus on one Z-slice and monitor recovery over time. This consisted of one prebleach image, bleaching at 100% laser power for 50-80 seconds, and a 2-stage time course. The first stage consisted of images taken every 5 seconds at scan speed 6 for 20 frames to track fast recovery. The second stage consisted of images taken every 30 seconds at scan speed 6 for 30 frames to track slower recovery. Laser power for tracking recovery was set below 5% to minimize unintentional photobleaching.

### FRAP Image Analysis and Model Fitting

The methodology as detailed in (Hiremath et al., 2024) was utilized to dissect the image data obtained from the FRAP experiments. Briefly, nuclei and their respective cytoplasmic compartments were delineated through segmentation and subsequently tracked across consecutive frames, enabling the extraction of both nuclear and cytoplasmic intensity profiles.

For one-component modeling, non-linear least squares fitting was employed to determine the kin (import rate) and kout (export rate) parameters. Briefly, the following equation was used to fit to the data:

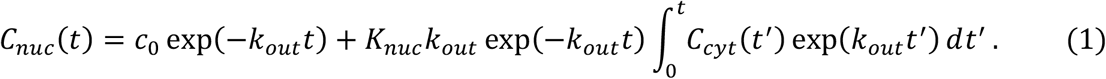

In Eqn (1), there are three adjustable parameters: *c*_0_, the estimated initial concentration after bleaching; *K*_*nuc*_ = *k*_*in*_/*k*_*out*_, the nuclear import/export equilibrium constant; and *k*_*out*_. In addition, *C*_*cyt*_ is the cytoplasmic intensity (measured from the intensity of the cytoplasmic domain bordering the bleached nucleus), and *C*_*nuc*_ is the intensity of the nucleus to be fit. The two-component model fit to the nuclear fluorescence was as follows:

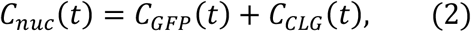

where “CLG” refers to Cact-LT/GFP. In this equation, the two components are each modifications of the one-component equation. For more details, see Supplementary Information.

### Live Embryo Imaging Compatible with Raster Image Correlation Spectroscopy (RICS)

Image acquisition compatible with RICS analysis was performed on embryos of 4 fly lines: 1×1x, 2x GFP, 2x Cact-LT, and the control. Each line was prepared as noted for whole body imaging in a previous section. A Zeiss LSM 900 confocal microscope was used to capture a 100-image time course of a single Z-slice at the embryo surface. Images were taken continuously at 1024×1024 pixels and a scan speed of 7, resulting in a pixel dwell time of 1 μs and a line time of 2.5 ms. Pixel size was 0.052 μm. Laser power was kept below 5% to minimize unintentional photobleaching.

### RICS analysis of Live Embryo Imaging and Model Fitting

RICS analysis was performed on the image acquisitions mentioned above as described in the literature (Al Asafen et al., 2018; Digman, Brown, et al., 2005; Digman, Sengupta, et al., 2005). Briefly, images were background subtracted, and the nuclear and cytoplasmic regions were segmented. Next, for each frame in the acquisition, immobile fractions of the image were removed using a sliding window average frame subtraction of five frames on either side of a given frame (Al Asafen et al., 2018; Digman, Brown, et al., 2005). Next, autocorrelation functions (ACFs) were found for each domain (nuclear and cytoplasmic) and each frame. The final ACF for an image acquisition and domain was the average of the ACFs for each frame.

The ACFs were then fit to either a one-component (Digman, Brown, et al., 2005; Digman, Sengupta, et al., 2005) or two-component model (Al Asafen et al., 2018). In the one component model, the theoretical ACF, *G*, was given as:

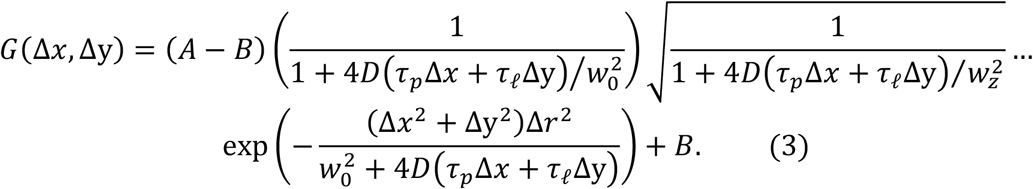

In this equation, there are three adjustable parameters: the amplitude of the ACF, *A*; the background levels, *B* (which are zero in theory, but the best fit could be non-zero in practice, so the parameter is adjustable for robustness of fit); and the diffusivity of the protein, *D*. The remaining variables are microscope parameters: *w*_0_ is a measure of the spatial extent of the excitation density in the *x y* plane (usually taken to be the radius of the first local minimum in the point spread function), and *w*_*z*_ is that along the axial (*Z*) direction; τ_*p*_ is the pixel dwell time, τ_*l*_ is the line time, and Δ_*r*_ is the pixel size.

In the two component model, a one component ACF with *D* = 4 μm^2^/s (*G*_-*cact*_; for Cact) is combined with a one-component ACF with *D* = 28 μm^2^/s (*G*_*GFP*,_ for free GFP), with a linear combination weight of *ϕ* and along the vertical scanning direction (Δ *y*) only (*i*.*e*., Δ*x* = 0):

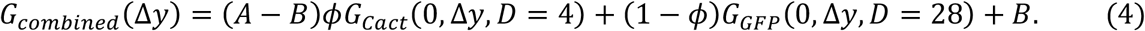

For more information on RICS analysis and fitting the ACFs, see the Supplementary Information and (Al Asafen et al., 2018).

Model fitting results were filtered for goodness-of-fit (*R*^2^) of greater than 0.85. Qualitative conclusions were unchanged by using lower thresholds.

## Results

### Cactus has a clear nucleocytoplasmic distribution

As a crucial player in embryo development, the Dl gradient has been quantified in live and fixed embryos (Al Asafen et al., 2020; Carrell et al., 2017; DeLotto et al., 2007; Liberman et al., 2009; Reeves et al., 2012). However, despite its central role in regulating the Dl gradient (Fig. 1A), quantification of Cact distribution is lacking. To visualize the dynamics of Cact in live embryos, we used CRISPR to tag the 3’ end of the endogenous *cact* locus with an LT that binds eGFP (Fig. 1B; see Methods; Bothma et al., 2018; Kirchhofer et al., 2010). With this construct, mature GFP molecules ubiquitously expressed in the embryo can bind the Cact-LT construct, allowing us to visualize Cact expression in both the nucleus and cytoplasm (Fig. 1C). We first analyzed live, whole mount embryos maternally expressing one copy of Cact-LT and one copy of eGFP (1×1x embryos; see Methods). These embryos also maternally expressed one copy of Histone-RFP.

**Figure 1:**
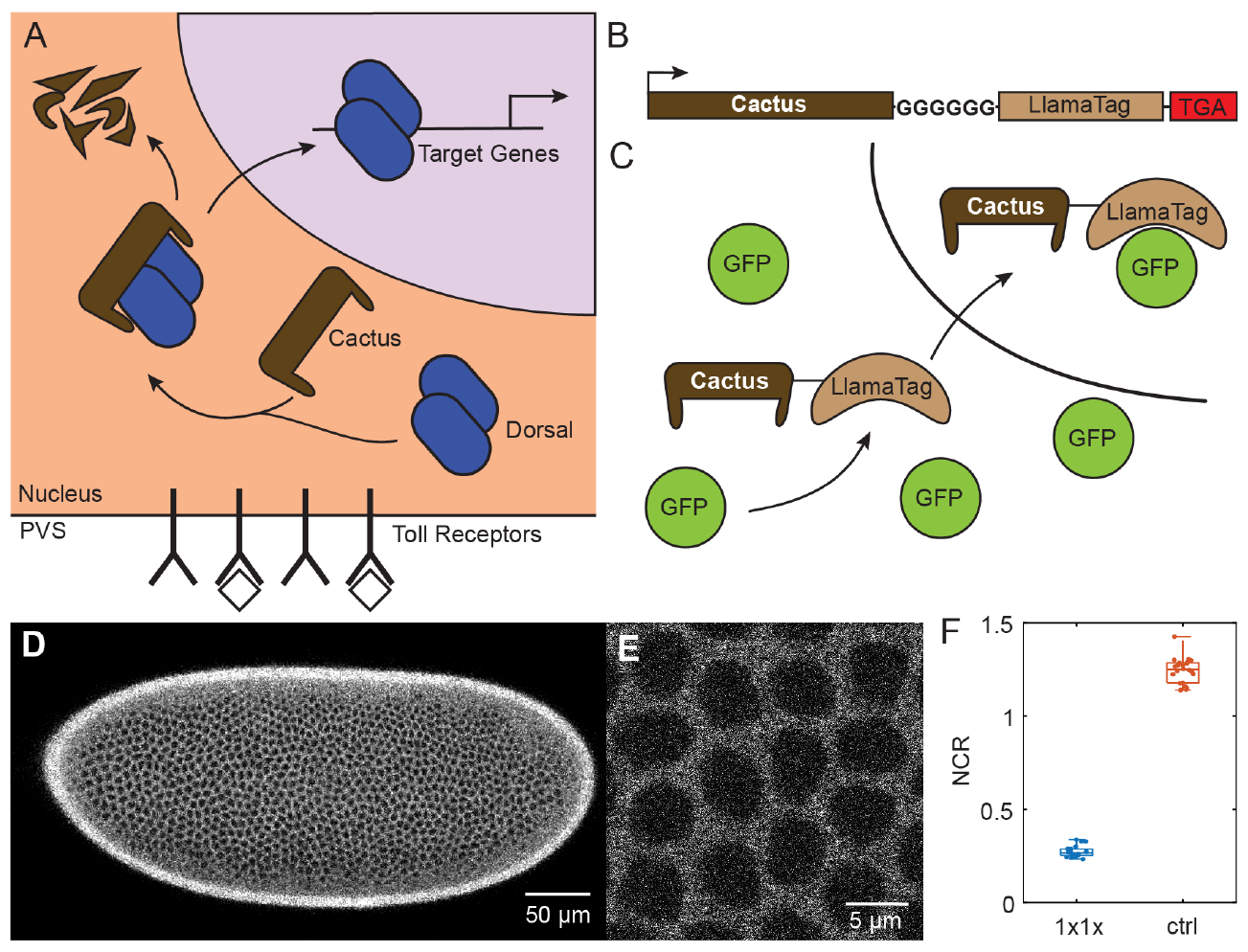
Cactus interactions and nucleocytoplasmic distribution. (A) A schematic showing the known role of Cact, where upon receiving a degradation signal via Toll, it dissociates from Dl, allowing Dl to enter the nucleus and interact with target genes. (B) A schematic of the Cact-LT fusion. The LlamaTag is tagged onto the 3’ end of the endogenous *cact* locus, connected by a glycine linker. (C) A schematic of Cact-LT’s interaction with GFP. (D) A whole-body image of 1×1x showing the cytoplasmic pattern. (D) A close-up image of 1×1x showing the cytoplasmic pattern. (F) A box plot of nucleocytoplasmic ratios for 1×1x and control embryos.

Imaging these embryos revealed a predominantly cytoplasmic pattern of eGFP fluorescence in blastoderm stage embryos (Fig. 1D, Movie S1). During nuclear cycles, this pattern disappeared as nuclear envelopes dissolved before being reestablished with the reformation of new nuclear envelopes (Movie S1). Control embryos expressing eGFP and Histone-RFP, but not Cact-LT, showed a slight nuclear localization of fluorescence (Fig. S1, Movie S2). These results suggest that eGFP is indeed associated with the LT fused to Cact, not just moving freely within the embryo, and that this association affects the distribution of GFP fluorescence.

We next quantified the nucleocytoplasmic distribution of eGFP fluorescence at higher resolution (Fig. 1E,F). We found the nuclear-to-cytoplasmic ratio (NCR) of fluorescence intensity was 0.27 ± 0.03 (Fig. 1E,F). In contrast, similar quantification of eGFP fluorescence in control embryos shows a weak nuclear localization (NCR = 1.24 ± 0.07; Figs. 1F and S1). Fixed 1×1x embryos showed a normal Dl gradient (Fig. S2A-C). Taken together, the presence of Cact-LT redistributes eGFP localization, which implies that Cact protein is predominantly cytoplasmic.

### Cactus protein is uniform along the DV axis in the blastoderm embryo

To determine whether there is a dorsal-ventral gradient of Cact expression, and whether Cact distribution changes in time, we imaged live embryos from nc 10 – 14 in a vertical orientation, which captures the full DV axis within the image frame (Fig. 2A-B, Movie S3; see Methods). Using computational image analysis methods (see Methods), we measured the nuclear and cytoplasmic intensities as functions of DV coordinate and time. We found no clear, persistent DV gradient in fluorescence intensity in either the nuclei or the cytoplasm (Fig. 2C).

**Figure 2:**
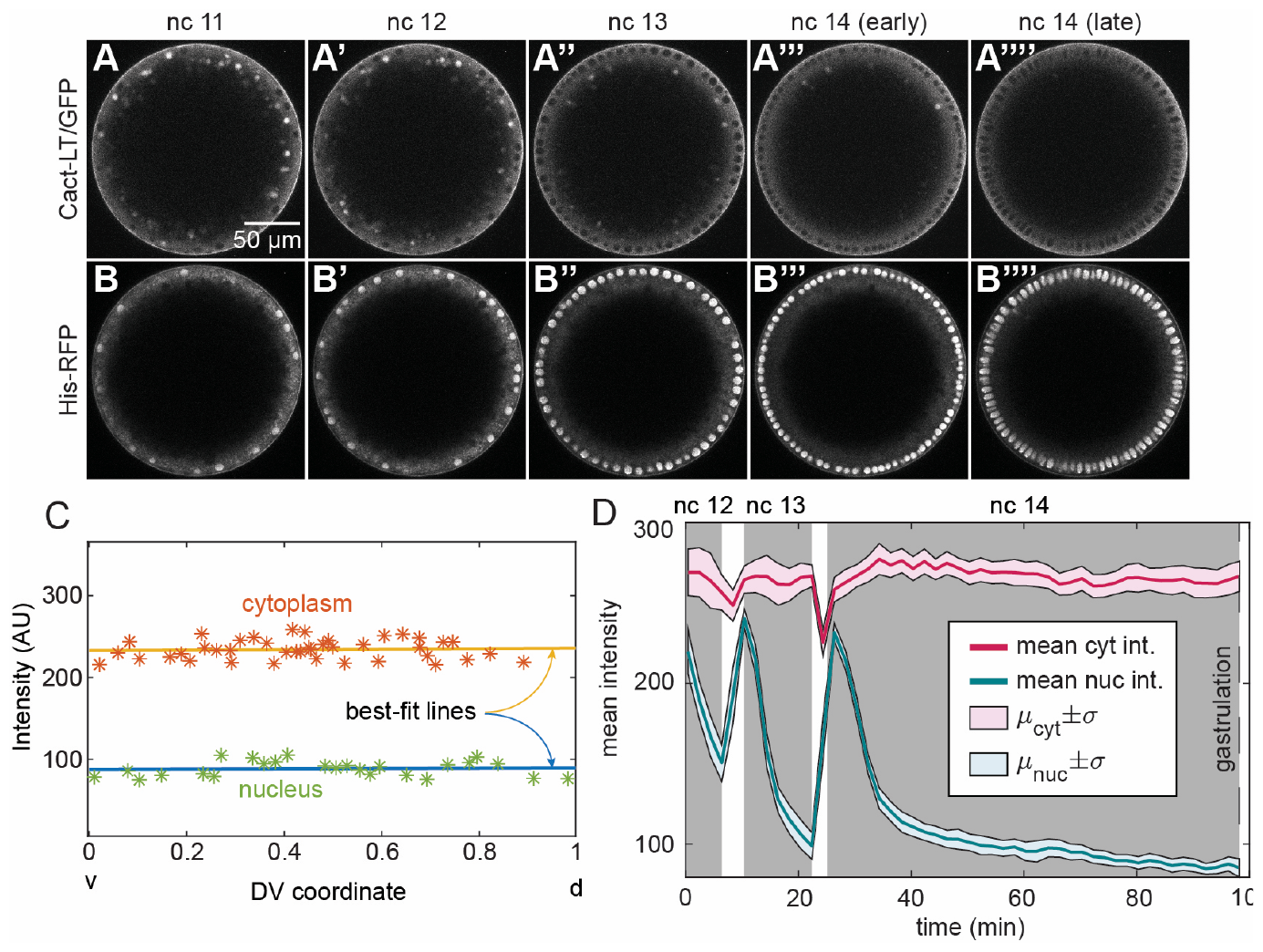
Cross-section analysis reveals uniform Cact gradient in the blastoderm embryo. (A-A’’’’) Images of Cact-LT/GFP expression from nc 11 to late nc 14. (B-B’’’’) Images of His-RFP expression, identifying nuclei, from nc 11 to nc 14. (C) Fluorescent intensity over the DV axis for the cytoplasm (orange) and nuclei (green) in cross-sectioned 1×1x embryos. (D) The mean fluorescent intensity over time for embryo cross-sections. The solid pink line represents the mean for the cytoplasm with the pink region representing the standard deviation. The solid blue line represents the mean for the nucleus over time with the blue region representing the standard deviation.

To quantify this, we treated each frame of the live imaging time course independently and tested whether there was a detectable slope to the intensity from the ventral to the dorsal side by performing simple linear least squares regression. We performed a *t*-test on the ventral-to-dorsal slope for each frame and found the number of frames with a p-value of less than 0.05 was consistent with this number occurring by chance.

Next, we measured the nuclear and cytoplasmic intensities over time. We found cytoplasmic intensities exhibited a relative stability across and during all nc 11-14 interphases (Fig. 2D). Transient decreases in cytoplasmic intensity were observed during mitosis, which would quickly recover to their baseline levels at the start of each interphase. In contrast, the study revealed distinct oscillations in nuclear intensities in which intensity would abruptly increase during mitosis, then slowly decrease during interphase, such that a steady state was reached only in nc 14 (Fig. 2D). This distinctive behavior of Cactus in the nuclei is similar to the pattern observed for fluorescently-tagged Dorsal in the dorsal-most nuclei (Reeves et al., 2012), which would suggest that, in both cases, it is Dl/Cact complex being observed (Carrell et al., 2017).

### Cactus exhibits nucleocytoplasmic shuttling

In the 1×1x embryos, the fluorescent intensity NCR, discerned from standard confocal imaging, measured to be roughly 0.3 after coming to steady state in the latter half of nc 14. This value suggested that Cact, widely known as a cytoplasmic tethering protein, is also present in the nuclei. However, due to the uniform expression of free GFP in these embryos, it is necessary to deconvolve the fluorescence arising from Cact-LT/GFP from that of free GFP fluorescence. To address this challenge, we performed fluorescent recovery after photobleaching (FRAP) to estimate the effect of fluorescence from the two sources. After photobleaching of a single nucleus (Fig. 3A; Movie S4), the fluorescence intensity exhibited a recovery trend over a time span of approximately 10-15 minutes (Fig. 3B). By fitting a model of Cact-LT/GFP nuclear import/export (transport) to the recovery curves (red curve in Fig. 3B), we estimated the nuclear import/export rate constants to be approximately *k*_*in*_ = 0.4 ± 0.4 and *k*_*out*_ = 1.1 ± 1.1 min^-1^, respectively (n = 16; Fig. 3C). The ratio was estimated to be *K*_*nuc*_ *k*_*in*_/*k*_*out*_ = 0.38 ± 0.04 (Fig. 3D), which is consistent with our NCR observations (Fig. 1F). In contrast, nuclei subjected to photobleaching in control embryos expressing solely free GFP, without Cact-LT (see Methods; Movie S5), exhibited a notably quicker recovery, with a time scale of about 10-20 seconds (*k*_*in*_ = 10.1 ± 3.9 min^-1^ and *k*_*out*_ = 8.9 ± 3.6 min^-1^; Fig. 3C; n = 12), and a ratio of *K*_*nuc*_ = 1.14 ± 0.04 (Fig. 3D), similar to our NCR measurements (Fig. 1F). These findings strongly support the presence of Cact-LT within the nucleus, as it significantly influences the nuclear transport rate constants of GFP.

**Figure 3.**
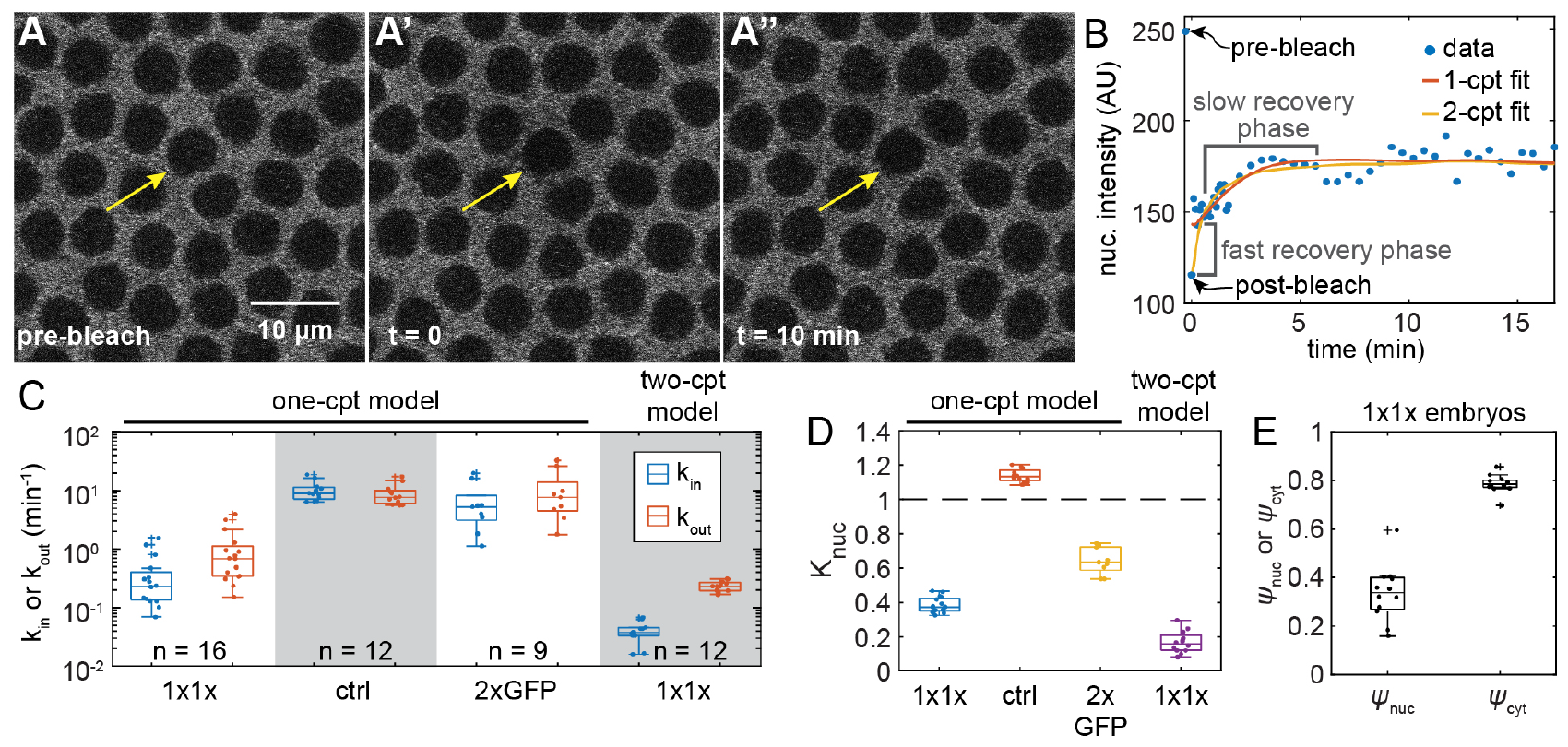
FRAP analysis provides evidence of Cact shuttling in and out of the nucleus. (A) Still frames of FRAP showing a nucleus prebleach [A], immediately after bleaching [A’], and 10 minutes post-bleach [A’’]. (B) Fluorescent recovery curves over time, fit using both the one-component and two-component model. (C) Boxplots of kin and kout for 1×1x, free GFP control, and 2x GFP using the one-component model, and kin and kout for 1×1x using the two-component model. (D) Boxplot of Knuc for 1×1x, free GFP control, and 2x GFP using the one-component model, followed by 1×1x using the two-component model. (E) Fraction of fluorescence attributable to Cact-LT/GFP in the nucleus and cytoplasm, inferred from the two-component model.

However, it is unlikely that 100% of the nuclear fluorescence stems from Cact-LT/GFP. Therefore, the FRAP recovery curves of 1×1x embryos likely represent a mixture of a fast-recovering free GFP and slow-recovering Cact-LT/GFP. To address this issue, we used a two-component model (yellow curve in Fig. 3B and Methods), which accounts for fluorescent recovery of both free GFP and Cact-LT/GFP. The model includes parameters for the nuclear transport rate constants of both species, as well as the fraction of fluorescence from Cact-LT in the nucleus and cytoplasm (*ψ*_*nuc*_ and *ψ*_*cyt*_, respectively), which correspond to the mol fraction of Cact. To ensure we did not overfit the model to the FRAP data in 1×1x embryos, we fixed the equilibrium constant for free GFP at 1.2 (Fig. 1F; see Supplementary Information). We found the two-component model predicted the nuclear transport rate constants of Cact-LT/GFP to be *k*_*in*_ = 0.04 ± 0.02 min^-1^ and *k*_*out*_ = 0.24 ± 0.05 min^-1^ (n = 12; Fig. 3C), the ratio of which was found to be *K*_*nuc*_ = 0.17 ± 0.06 (Fig. 3D). The model also predicted that roughly one third of nuclear fluorescence and 80% of cytoplasmic fluorescence stems from Cact-LT/GFP (Fig. 3E; *ψ*_*nuc*_ = 0.34 ± 0.11; *ψ*_*cyt*_ = 0.79 ± 0.04).

To further test whether the recovery is a mixture of free GFP and Cact-LT/GFP, we varied the dosage of GFP. In particular, higher total GFP levels should result in faster apparent recovery rates in the presence of Cact-LT, because a greater fraction of the mixture should be free GFP. Therefore, we performed FRAP in 2x GFP embryos and, using the one-component model for FRAP recovery, we found the apparent nuclear transport rate constants to be *k*_*in*_ = 7.0 ± 6.4 and *k*_*out*_ = 11.1 ± 10.7 min^-1^ (n = 9; Fig. 3C; Movie S6). Because the recovery rates are driven mostly by free GFP, the estimated nuclear transport rate constants are on the same order of magnitude as those in control embryos (free GFP only). For the same reason, we were not able use the two-component model to reliably deconvolve the relatively minor effect of Cact-LT/GFP from free GFP. However, it should be noted that the ratio of *K*_*nuc*_ = *k*_*in*_/*k*_*out*_ is still less than one (0.64 ± 0.08) due to the presence of Cact-LT (Fig. 3D).

### Cactus represents roughly half of the nuclear fluorescence

While FRAP provided clear evidence of nuclear Cact, and the two-component model provided an estimate of the fraction of fluorescence from Cact-LT/GFP (*ψ*_*nuc*_ and *ψ*_*cyt*_), some of the parameter estimates had high uncertainty. Therefore, we sought an independent estimate of *ψ*_*nuc*_. To this end, we used raster image correlation spectroscopy (RICS; Al Asafen et al., 2018; Brown et al., 2008; Digman, Brown, et al., 2005; Digman, Sengupta, et al., 2005), a derivative fluorescence correlation spectroscopy (Elson & Magde, 1974). RICS analysis entails calculating correlations in statistical fluctuations between closely timed voxels to measure the absolute concentration and mobility of multiple fluorescent species. The analysis is performed on a time series of high magnification laser scanning confocal images (Fig. 4A-C, Movie S7), and results in a two-dimensional autocorrelation function (ACF) that is a function of pixel shifts in the fast and slow scanning directions of the confocal microscope (Al Asafen et al., 2018). However, for the purposes of estimating the diffusivity of fluorescent molecules, the slow scanning direction (y-direction in the image) is the key component of the ACF (diamonds and circles in Fig. 4D). A theoretical ACF for a diffusible fluorescent molecule is then fit to the ACF calculated from image data to infer the diffusivity and concentration of the fluorescent molecule (Fig. 4D). RICS analysis can focus on different domains in the image, such as calculating the diffusivity in the nucleus and cytoplasm separately (see Methods; Al Asafen et al., 2018). Furthermore, if multiple fluorescent species are present that have substantially different diffusivities, RICS analysis can infer the relative abundances of the two species (Al Asafen et al., 2018).

**Figure 4.**
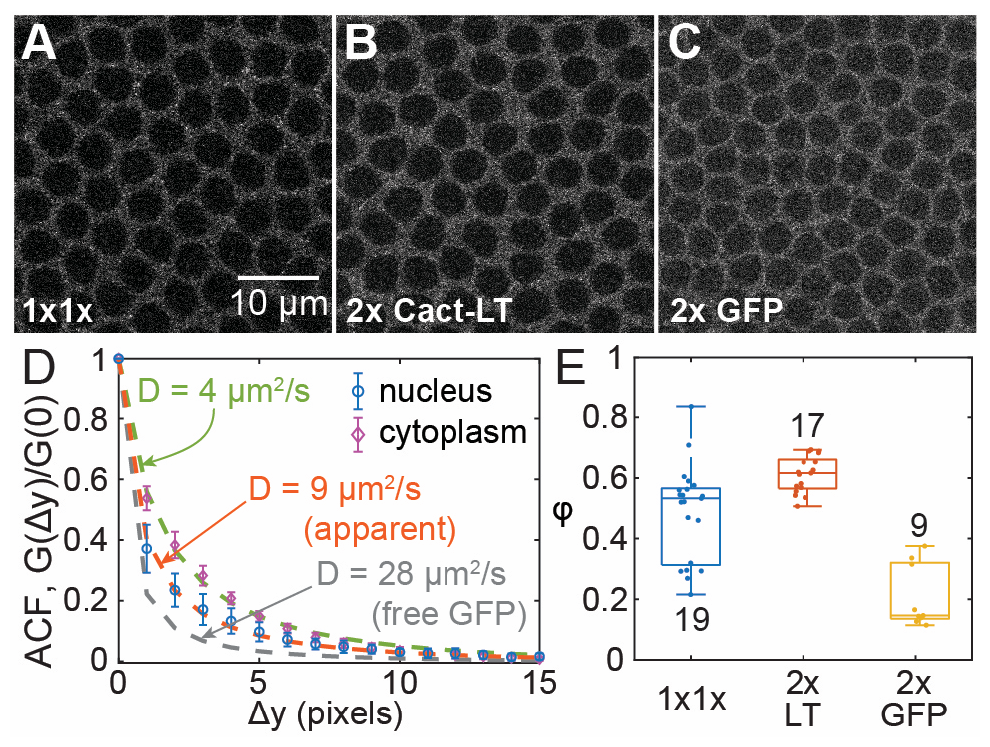
RICS analysis provides evidence of Cact-LT nuclear fraction. (A) Still frame of 1×1x embryo imaged for RICS analysis. (B) Still frame of 2x Cact-LT embryo imaged for RICS analysis. (C) Still frame of 2x GFP embryo imaged for RICS analysis. (D) Average nuclear (blue circles) and cytoplasmic (purple diamonds) ACFs for n = 16 1×1x embryos (normalized). Calculated diffusivity rates for cytoplasmic Cact-LT (fit line in green), nuclear Cact-LT (fit line in pink). For comparison, an ACF corresponding to D = 28 μm^2^/s is plotted in gray, consistent with free GFP movement. Errorbars: standard deviation. (E) Boxplot of mol fraction of Cact-LT in nuclei for 1×1x, 2x Cact-LT, and 2x GFP. Sample side displayed on plot.

In the cytoplasm, where the GFP fluorescence is likely to be predominantly Cact-LT/GFP, the diffusivity was D = 3.9 ± 0.5 μm^2^/s (n = 19) (Fig. 4D). However, in the nucleus, the apparent diffusivity was significantly higher (D = 9 ± 3 μm^2^/s; n = 23). In control embryos, the diffusivity of free GFP in the nuclei was D = 28 ± 1 μm^2^/s (n = 22; Fig. 4D): roughly the same as a protein diffusing in water. This suggests that the intermediate apparent diffusivity of the fluorescent molecules in the nucleus is a combination of Cact-LT/GFP, which diffuses slowly, and free GFP, which has high diffusion. Assuming that, in these embryos, free GFP molecules still have a diffusivity of 28 μm^2^/s, and that Cact-LT/GFP in the nucleus has the same diffusivity as measured in the cytoplasm (∽ 4 μm^2^/s), we fit a linear combination of slow and fast diffusing ACF models to the ACF data, with *ϕ* as the linear combination weight parameter (see Methods and Al Asafen et al., 2018). The larger *ϕ* is, the more that Cact-LT/GFP contributes to the nuclear GFP fluorescence. Furthermore, if the two species have the same brightness, then *ϕ* represents the mol fraction of Cact-LT/GFP out of all GFP-containing species:

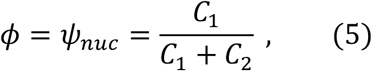

where *C*_1_ is the concentration of Cact-LT in the nucleus, and *C*_2_ is the concentration of free GFP in the nucleus.

After fitting the combined ACF to the ACF derived from the nuclear fluorescence fluctuation data, we found *ϕ* = 0.47 ± 0.16 (Fig. 4E; see Methods). These numbers are consistent, albeit slightly higher, than those estimated from the two-component FRAP model. Therefore, a significant fraction of fluorescence in the nucleus stems from Cact-LT, which corresponds to roughly half of the GFP if the two species have the same brightness.

On the other hand, the LlamaTag fusion protein used in this study and others (Bothma et al., 2018) has been reported to increase GFP fluorescence by 1.5-fold (Kirchhofer et al., 2010). In that case, *ϕ* is defined as:

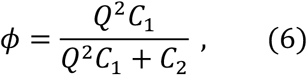

where *Q* is the fold increase of GFP brightness upon binding the llamatag. If *ϕ* = 0.5 and *Q* = 1.5, then the mol fraction of Cact-LT/GFP out of all GFP-containing species is

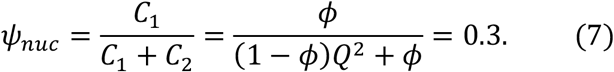

If *ϕ* is indeed a measure of the relative amount of Cact-LT/GFP is present in the nucleus, it should be an increasing function of Cact-LT dosage and a decreasing function of free GFP dosage. Therefore, we measured this parameter in embryos maternally expressing two copies of Cact-LT and one copy of free GFP (2x Cact-LT; Movie S8) or one copy of Cact-LT and two copies of free GFP (2x GFP; Movie S9). As expected, in the 2x Cact-LT embryos, *ϕ* increased to 0.60 ± 0.06, and in the 2x GFP embryos, it decreased to 0.18 ± 0.10 (Fig. 4E). We conclude that *ϕ* is an effective measure of the presence of Cact-LT in the nucleus and that a substantial fraction of GFP in the nucleus is bound by Cact-LT. In particular, from the RICS and FRAP analyses in 1×1x embryos, the mol fraction of GFP in the nucleus bound by Cact-LT is 0.3-0.5.

### Concentration of Cactus and the binding affinity of GFP to the llamatag

Given the above measurements of Cact biophysical parameters using RICS and FRAP, we surmised there may be sufficient constraints to infer the remaining unknown system parameters, such as the concentration of Cact and the dissociation constant for the Cact-LT/GFP interaction. Therefore, we developed a material balance model of Cact-LT/GFP dynamics and constrained it with our quantitative measurements of the nuclear import/export rates, mol fraction of Cact-LT/GFP and the NCR of intensities in 1×1x, 2xGFP, 2xLT, and control embryos, as well as an estimate of the total GFP concentration in the embryo (see Supplementary Information). We used an evolutionary optimization strategy, ISRES+ (Bandodkar et al., 2023; see Supplementary Information), which results in distributions of inferred parameters (Fig. 5). In particular, we estimate forward binding rate constant between Cact-LT and GFP to be kon = 0.03 ± 0.01 nM^-1^min^-1^ (Fig. 5A) and the dissociation constant to be KD = 21.80 ± 5.11 nM (Fig. 5B). These estimates characterize the interactions between the LT and GFP, and thus, could be generally used in interpretations of experiments involving this LT (Bothma et al., 2018; Garcia et al., 2020). Furthermore, the total Cact concentration (averaged between nuclear and cytoplasmic compartments) is estimated to be = 133 ± 31 nM, which implies a nuclear concentration of 15 ± 4 nM and a cytoplasmic concentration of 176 ± 40 nM (Fig. 5C). The NCR of these concentrations (roughly 0.1) is consistent with our two-component FRAP model (Fig. 3D). These estimates of Cact biophysical parameters can help constrain predictive models of the Dorsal gradient (Ambrosi et al., 2014; Barros et al., 2021; Carrell et al., 2017; Kanodia et al., 2009; O’Connell & Reeves, 2015).

**Figure 5:**
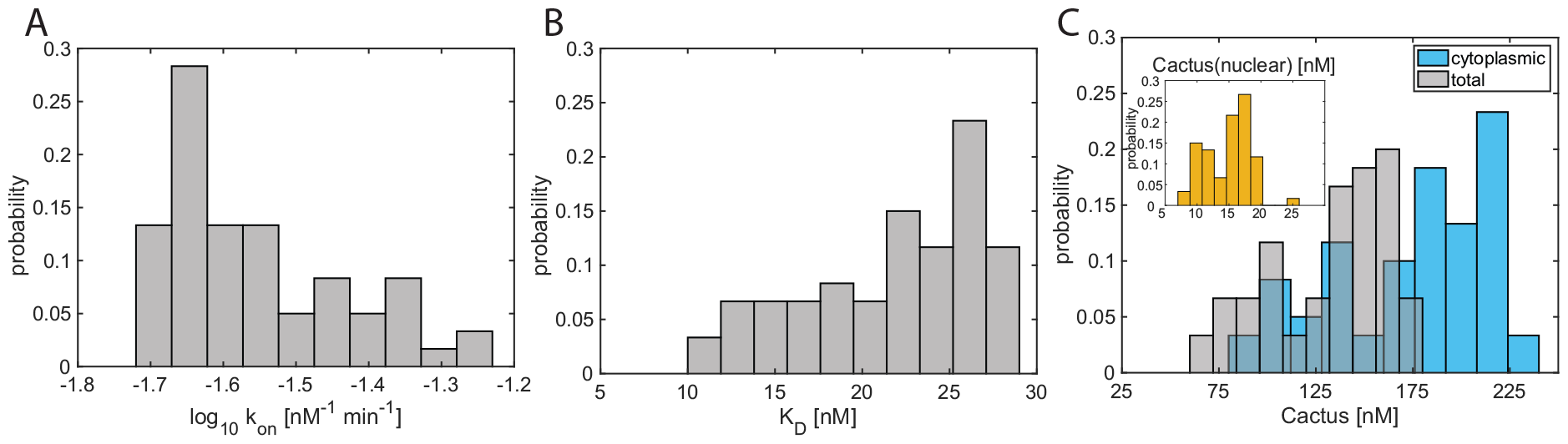
Model inferences. Histograms of the distributions of kon (A), KD (B), and Cact concentrations (C).

## Discussion

In this paper, we used advanced confocal imaging techniques to measure the spatiotemporal distribution of a LlamaTag-fused Cact (Cact-LT) bound to GFP in the *Drosophila* blastoderm embryo. We also used mathematical modeling to infer biophysical properties from the experimental data, such as total Cact concentration in the embryo and the dissociation constant of the LlamaTag and GFP. Our results provide new insights into how Cact regulates the Dorsal/NF-κB gradient in blastoderm-stage *Drosophila* embryos.

Our imaging results show that, while GFP fluorescence is predominantly cytoplasmic, a non-negligible amount of fluorescence resides in the nuclei, such that, at steady state, the nuclear-to-cytoplasmic ratio (NCR) of fluorescence intensity was roughly 1/3. However, GFP is expressed ubiquitously, and only a fraction of the GFP is bound by Cact-LT. Therefore, using two independent methods (FRAP and RICS), we measured the fraction of nuclear fluorescence that stems from Cact-LT and found it to be roughly one 0.3-0.5. It is believed that most of the Cact present in the embryo is bound to Dl because, in embryos with classical knockout alleles of Dl, Cact levels become undetectable (Govind et al., 1993; Kidd, 1992; Whalen & Steward, 1993). From this, we surmise that Dl/Cact complex is the primary Cact-containing species localized to the nuclei. Therefore, the nuclear import and export rates of Cact-LT/GFP (0.04 and 0.24 min^-1^, respectively) are likely to be measurements of Dl/Cact complex nuclear transport.

In vertebrates, the Cact homolog IκBα is known to enter the nucleus to regulate NF-κB transcription factors (Arenzana-Seisdedos et al., 1995, 1997; Kearns et al., 2006). In contrast, *Drosophila* Cact has been presumed to be purely cytoplasmic (Belvin et al., 1995; Bergmann et al., 1996; Roth et al., 1991; Whalen & Steward, 1993). Our previous studies have challenged this view (Al Asafen et al., 2018, 2020; O’Connell & Reeves, 2015), providing indirect evidence that Cact is in the nucleus. However, due to the high turnover rate of Cact protein, direct in situ visualization of Cact protein (in the nucleus or elsewhere) has been sparse (Belvin et al., 1995; Bergmann et al., 1996; Whalen & Steward, 1993). Therefore, our direct visualization of Cact in the nucleus has major implications for our understanding of the Dl gradient. In particular, it implies that nuclear fluorescence measurements of the Dl gradient include both free (active) and Cactbound (inactive) Dl. In other words, the presence of Dl/Cact complex in the nucleus obfuscates our measurements of the true Dl activity gradient. Thus, future quantification of the Dl activity gradient (free Dl alone) must include methods to deconvolve inactive Dl/Cact complex from the total Dl gradient (free Dl + Dl/Cact complex), either with accurate computational models (Al Asafen et al., 2020; O’Connell & Reeves, 2015), aided by quantitative measurements such as those in this paper, or with experimental methods that can measure protein-protein complexes, such as cross-correlation RICS (Digman et al., 2009).

Our imaging of full dorsal-ventral optical cross sections during the blastoderm stage found that there is no discernible dorsal/ventral polarity to the Cact distribution, either in the cytoplasm or in the nuclei. This finding is consistent with our previous modeling of the Dl gradient, in which the Dl/Cact complex distribution is roughly flat across the embryo (Carrell et al., 2017; O’Connell & Reeves, 2015). While this result may seem surprising given that Toll-dependent degradation of Cact is localized on the ventral side of the embryo, we have previously shown that ventrally-directed diffusion of Dl/Cact complex (i.e., “shuttling” of Dl) is sufficient to flatten the gradient of Dl/Cact complex (Carrell et al., 2017).

Even with a flat gradient of Dl/Cact complex, deconvolving inactive Dl/Cact complex from total Dl results in a Dl activity gradient that carries more positional information than the total Dl gradient measured by standard fluorescence (O’Connell & Reeves, 2015). This is because subtracting out the contribution from Dl/Cact complex in the nuclei results in a Dl activity gradient that decays to near zero, which dramatically increases the dynamic range of the Dl gradient to multiple logs (O’Connell & Reeves, 2015). This results in a gradient that is more robust to intrinsic noise and to variations in overall Dl concentration (Al Asafen et al., 2020; O’Connell & Reeves, 2015). Ultimately, without deconvolution, the expression patterns of Dl target genes cannot be properly explained by fluorescence measurements of the gradient (Al Asafen et al., 2020; O’Connell & Reeves, 2015; Reeves et al., 2012).

In conclusion, through use of the novel LlamaTag technology, our study uncovered new insights into the regulation of the Dl gradient and its target genes by providing direct experimental evidence of the presence of Cact in the nuclei of blastoderm-stage embryos. Furthermore, our work highlights the importance of synergizing experimental methods, including advanced imaging techniques, with mathematical modeling to advance our understanding of biological systems.

