## Supplemental Information for "Spatiotemporal dynamics of NF-κB/Dorsal inhibitor IκBα/Cactus in *Drosophila* blastoderm embryos"

### Supplementary Information for "Spatiotemporal dynamics of NF-κB/Dorsal inhibitor IκBα/Cactus in *Drosophila* blastoderm embryos"

Allison E. Schloop[a], Sharva Hiremath[b], Razeen Shaikh[c], Cranos M. Williams[b], and Gregory T. Reeves[c,d,1]

[a] Graduate Program in Genetics, North Carolina State University, Raleigh, NC 27606

[b] Department of Electrical and Computer Engineering, North Carolina State University, Raleigh, NC 27606

[c] Department of Chemical Engineering, Texas A&M University, College Station, TX 77843

[d] Faculty in Genetics and Genomics, Texas A&M University, College Station, TX 77843

#### Generating the averaged plot for the Cact embryo cross sections

The following steps were used to generate the averaged plot for Cact embryo cross section:

1) Scale and offset: Detect the nc13 and nc14 peaks (in nuclear intensities) for the different embryos by evaluating their respective intensities and identifying the two highest peaks (indicates start of nc13 and nc14). After detecting the peaks, aligned the curves such that the 13 and nc14 peaks are in line for the 4 embryos by calculating the scale and offset for transformation of the intensities in the time axis.

   Since different embryo data might have been collected from different points in the nuclear cycles, this step ensures that all the intensities are at equivalent time points during comparison.

   Finally, the amplitudes of the intensity curves are scaled such that the peaks have the same amplitude. This eliminates the discrepancies that arises from collecting data at different depths from embryos.

2) Interpolations: Since we offset the time-axis for some curves, we used interpolation to ensure that the intensity values of different embryos are all along the same time points before calculating the mean. Additionally, we extrapolated the intensities for curves whose time was offset (to ensure that all curves have the same time axis).

3) Mean: Calculated the mean for the new interpolated nuclear and cytoplasmic curves

4) Error propagation: Used error propagation to calculate the new standard deviation for nuclear and cytoplasmic intensities.

#### Model fitting of FRAP data – one component

The nuclear GFP intensity, $C_{nuc}$, is modeled by the following differential equation:

$$\frac{dC_{nuc}}{dt} = k_{in}C_{cyt}(t) - k_{out}C_{nuc},$$

where $k_{out}$ is the nuclear export rate constant, $k_{in}$ is the nuclear import rate constant, and $C_{cyt}(t)$ is the measured intensity of the cytoplasm. This equation can be solved analytically to yield:

$$C_{nuc}(t) = c_0 \exp(-k_{out}t) + k_{in} \exp(-k_{out}t) \int_0^t C_{cyt}(t') \exp(k_{out}t')\, dt',$$

where $c_0$ is the estimate of the initial concentration after bleaching. Because $C_{cyt}$ is defined at discrete time points, we assumed $C_{cyt}(t')$ to be a continuous, piecewise linear function between the data points. In other words, between any two time points $i$ and $i + 1$:

$$C_{cyt}(t) = \frac{\Delta C_{cyt,i}}{\Delta t_i}(t - t_i) + C_{cyt}(t_i),$$

where $\Delta C_{cyt,i} = C_{cyt}(T_{i+1}) - C_{cyt}(t_i)$, $\Delta t_i = T_{i+1} - t_i$, and $T_{i+1}$ is the minimum of $\{t_{i+1}, t\}$.

Substituting the above approximation for $C_{cyt}(t)$ into the solution to the differential equation allowed us to evaluate the integral analytically, which in turn allowed us to avoid numerical issues with evaluating this integral, especially if $k_{out}$ is large:

$$C_{nuc}(t) = c_0 \exp(-k_{out}t) + \cdots$$

$$k_{in} \exp(-k_{out}t) \sum_{i=1}^{n_t} \int_{t_i}^{T_{i+1}} \left[\frac{\Delta C_{cyt,i}}{\Delta t_i}(t' - t_i) + C_{cyt}(t_i)\right] \exp(k_{out}t')\, dt'$$

where $n_t$ is the number of points of data in the time interval $[0, t)$. Therefore:

$$C_{nuc}(t) = c_0 \exp(-k_{out}t) + \cdots$$

$$k_{in} \exp(-k_{out}t) \sum_{i=1}^{n_t} \int_{t_i}^{T_{i+1}} \left[\frac{\Delta C_{cyt,i}}{\Delta t_i} t' \exp(k_{out}t') + \left(C_{cyt}(t_i) - \frac{\Delta C_{cyt,i}}{\Delta t_i} t_i\right) \exp(k_{out}t')\right] dt'$$

$$= c_0 \exp(-k_{out}t) + \cdots$$

$$\frac{k_{in}}{k_{out}} \exp(-k_{out}t) \sum_{i=1}^{n_t} \left[\frac{\Delta C_{cyt,i}}{k_{out}\Delta t_i} [\exp(k_{out}T_{i+1})\,(k_{out}T_{i+1} - 1) - \exp(k_{out}t_i)\,(k_{out}t_i - 1)] + \left(C_{cyt}(t_i) - \frac{\Delta C_{cyt,i}}{\Delta t_i} t_i\right) [\exp(k_{out}T_{i+1}) - \exp(k_{out}t_i)]\right]$$

$$= c_0 \exp(-k_{out}t) + \cdots$$

$$K_{nuc} \sum_{i=1}^{n_t} \left[\frac{\Delta C_{cyt,i}}{k_{out}\Delta t_i} \left[\exp\left(-k_{out}(t - T_{i+1})\right)(k_{out}T_{i+1} - 1) - \exp\left(-k_{out}(t - t_i)\right)(k_{out}t_i - 1)\right] + \left(C_{cyt}(t_i) - \frac{\Delta C_{cyt,i}}{\Delta t_i} t_i\right) \left[\exp\left(-k_{out}(t - T_{i+1})\right) - \exp\left(-k_{out}(t - t_i)\right)\right]\right],$$

where $K_{nuc} = k_{in}/k_{out}$ is the nuclear import/export equilibrium constant.

**Model fitting of FRAP data – two component**

The two-component model fit to the nuclear fluorescence was as follows:

$$C_{nuc}(t) = C_{GFP}(t) + C_{CLG}(t),$$

where "CLG" refers to Cact-LT/GFP. We assumed the two components do not interact; that is, the association/dissociation of GFP from the llamatag did not appreciably affect the dynamics of the two species. Under such a decoupling assumption, the two components are simply modifications of the one-component equation:

$$C_{GFP}(t) = (1-\psi_{nuc})c_0 \exp(-k_{outG}t) + K_{nucG}k_{outG}\exp(-k_{outG}t)\int_0^t (1-\psi_{cyt})C_{cyt}(t')\exp(k_{outG}t')\,dt',$$

and

$$C_{CLG}(t) = \psi_{nuc}c_0 \exp(-k_{out}t) + K_{nuc}k_{out}\exp(-k_{out}t)\int_0^t \psi_{cyt}C_{cyt}(t')\exp(k_{out}t')\,dt',$$

where $\psi_{nuc}$ and $\psi_{cyt}$ are the mol fractions of Cact-LT/GFP (normalized to total GFP) in the nucleus and cytoplasm, respectively; $K_{nucG}$ and $K_{nuc}$ are the nuclear import/export equilibrium constants for free GFP and Cact-LT/GFP, respectively; and $k_{outG}$ and $k_{out}$ are the nuclear export rate constants of free GFP and Cact-LT/GFP, respectively. To avoid overfitting, $K_{nucG}$ was fixed at 1.2 (Fig. 3D); $c_0$ was fixed at the initial post-bleach intensity of the nucleus; and the following constraints, derived from material balances, were enacted for $\psi_{cyt}$ and $K_{nuc}$:

$$\psi_{cyt} = 1-(1-\psi_{nuc})\frac{NCR}{K_{nucG}},$$

$$K_{nuc} = \frac{\psi_{nuc}}{\psi_{cyt}}NCR,$$

where $NCR$ is the ratio of the steady state intensity of the bleached nucleus to the steady state intensity of the associated cytoplasm. In practice, these steady state intensities were taken as the average of all time points for $t > 6$ min post-bleach. These constraints left three adjustable parameters: $k_{out}$, $k_{outG}$, and $\psi_{nuc}$.

**Model fitting of RICS data**

In the one component model, the theoretical ACF, $G$, was given as:

$$G(\Delta x, \Delta y) = (A-B)\left(\frac{1}{1+4D(\tau_p\Delta x+\tau_\ell\Delta y)/w_0^2}\right)\sqrt{\frac{1}{1+4D(\tau_p\Delta x+\tau_\ell\Delta y)/w_z^2}}\cdots$$

$$\exp\left(-\frac{(\Delta x^2+\Delta y^2)\Delta r^2}{w_0^2+4D(\tau_p\Delta x+\tau_\ell\Delta y)}\right)+B.$$

In this equation, there are three adjustable parameters: the amplitude of the ACF, $A$; the background levels, $B$ (which are zero in theory, but the best fit could be non-zero in practice, so the parameter is adjustable for robustness of fit); and the diffusivity of the protein, $D$. The remaining variables are microscope parameters: $w_0$ is a measure of the spatial extent of the excitation density in the $xy$ plane (usually taken to be the radius of the first local minimum in the point spread function), and $w_z$ is that along the axial ($z$) direction; $\tau_p$ is the pixel dwell time, $\tau_\ell$ is the line time, and $\Delta r$ is the pixel size. In the slow direction, $\Delta x = 0$, and the ACF reduces to:

$$G(0, \Delta y) = (A - B)\left(\frac{1}{1 + 4D\tau_\ell \Delta y / w_0^2}\right)\sqrt{\frac{1}{1 + 4D\tau_\ell \Delta y / w_z^2}} \exp\left(-\frac{\Delta y^2 \Delta r^2}{w_0^2 + 4D\tau_\ell \Delta y}\right) + B.$$

In the two component model, a one component ACF with $D = 4$ μm$^2$/s (for Cact) is combined with a one-component ACF with $D = 28$ μm$^2$/s (for free GFP), with a linear combination weight of $\phi$:

$$G_{combined}(\Delta y) = (A - B)\phi G_{Cact}(0, \Delta y, D = 4) + (1 - \phi) G_{GFP}(0, \Delta y, D = 28) + B$$

**Material balance model:**

The steady-state mathematical model of the Cact-LT/GFP interaction has two compartments, nucleus and cytoplasm; and six species Cactus-LT (C), GFP (G) and the Cact-LT/GFP complex (CLG) in the nucleus and cytoplasm, denoted with the subscript 'n' and 'c', respectively. The model has five parameters: import rate ($k_{in}$), export rate ($k_{out}$), rate of formation of Cact-LT/GFP ($k_{on}$), binding affinity of Cact-LT to GFP ($K_D$), total Cactus concentration ($Cact_{tot}$).

$$\frac{dC_n}{dt} = k_{in} C_c - k_{out} C_n - k_{on} K_{nucG} G_c C_n + k_{off} CLG_n$$

$$\frac{dCLG_n}{dt} = k_{in} CLG_c - k_{out} CLG_n + k_{on} K_{nucG} G_c C_n - k_{off} CLG_n$$

$$\frac{dG_c}{dt} = -\frac{k_{on}}{1 + K_{nucG}}\left[(C_n - K_D CLG_n) + (C_c G_c - K_D CLG_c)\right]$$

$$0 = \frac{V_c}{V_{tot}}(CLG_c + C_c) + \frac{V_n}{V_{tot}}(CLG_n + C_n) - Cact_{tot}$$

$$0 = \frac{V_c}{V_{tot}}(CLG_c + G_c) + \frac{V_n}{V_{tot}}(CLG_n + G_n) - GFP_{tot}$$

We implemented Improved Stochastic Ranking Evolutionary Strategy *plus* (ISRES+; Bandodkar et al., 2023) to estimate the five model parameters. The model was fit to ϕ and NCRI for 1x1x, 2xGFP and 2xLT. The objective was to minimize:

$$f(params) = \sum_{i=1}^{3}\left(\frac{\frac{CLG_{n,i}}{G_{n,i} + CLG_{n,i}} - \phi_i}{\sigma_{\phi i}}\right)^2 + \sum_{i=1}^{3}\left(\frac{\frac{CLG_{n,i} + G_{n,i}}{CLG_{c,i} + G_{c,i}} - NCR_{I,i}}{\sigma_{NCRi}}\right)^2$$

where $i \in \{1x1x, 2xGFP, 2xLT\}$

ISRES+ is a global optimization algorithm which uses a (μ-λ) evolutionary strategy and employs two gradients-based strategies Linstep and Newton-step to efficiently estimate model parameters.

The model was run for 1000 generations with 150 individuals, with a recombination rate of 0.85. The contribution of Linstep and Newton-step were 2 and 1, respectively. The complete description of hyperparameters supplied to ISRES+ is in the Table S3.

The system of equations was solved to steady state computed with fsolve in MATLAB, with ISRES+ by randomly choosing a value of $GFP_{tot}$ and $K_{nucG}$ from a uniform distribution generated from their minimum and maximum estimates from RICS in 60 independent runs. fsolve's trust-region algorithm was used with a function tolerance of 1e-20 and optimality tolerance of 1e-20 and the initial conditions were randomly chosen for each independent simulation. Additionally, the upper and lower bounds of import and export rates were constrained based on experimental estimates. The value of $GFP_{tot}$ was doubled to simulate $\phi$ and NCRI for 2xGFP and the parameter $Cact_{tot}$ was doubled to simulate 2xLT. The model fits are as shown in Figure S4.

### Supplemental Figures and Tables

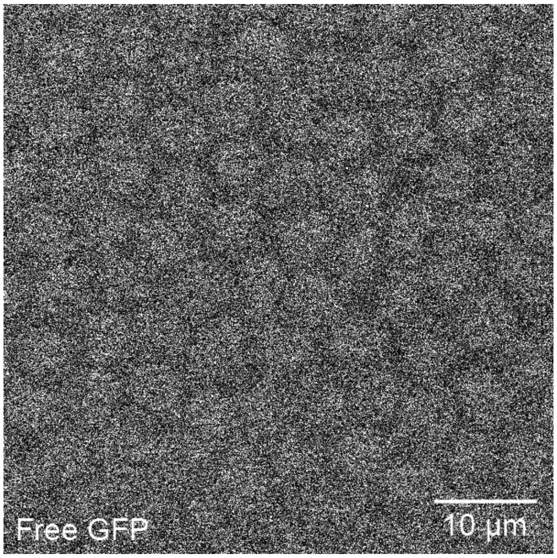

**Figure S1: Nucleocytoplasmic distribution of GFP in free GFP control embryo.**

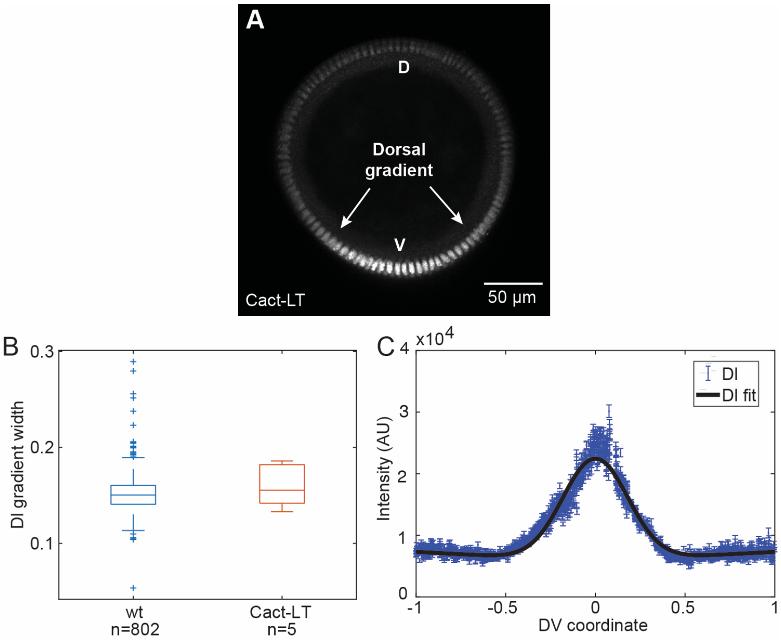

**Figure S2: Dorsal gradient of a fixed 1x1x embryo.** (A) Image of a 1x1x fixed embryo with Dl gradient marked. (B) Boxplot comparison of WT DV axis width versus 1x1x DV axis width showing no significant difference (p=0.33) (C) Fluorescent intensity over the [illegible]

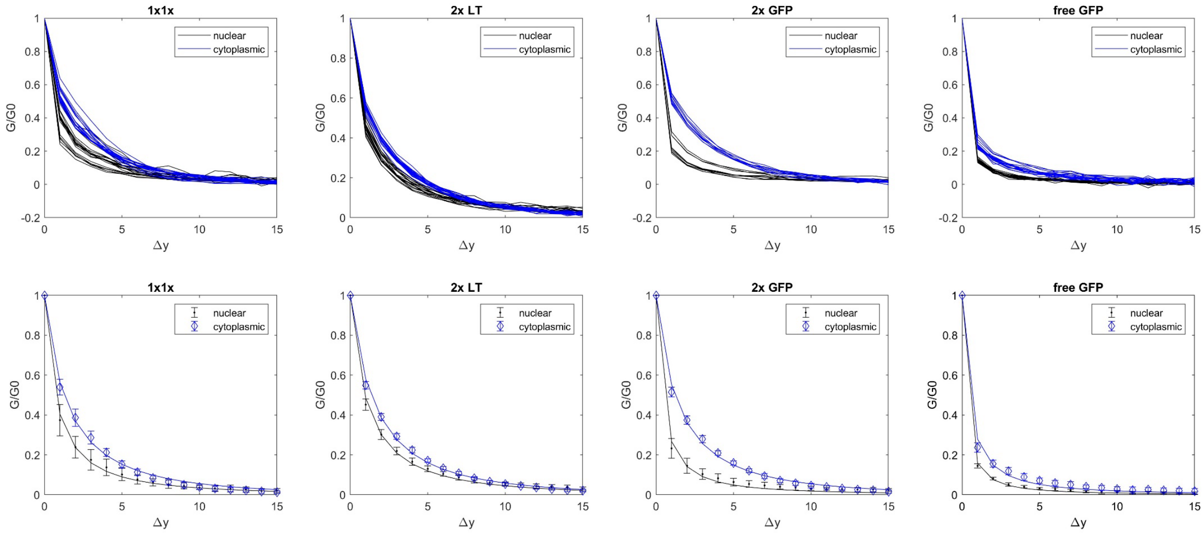

**Figure S3: RICS autocorrelation functions. Top row**: plot of all RICS ACFs along $\Delta y$ (*i.e.,* $\Delta x = 0$) for each genotype (in separate columns). **Bottom row**: plot of average and standard deviation of each pixel shift (symbol with errorbar) for all RICS ACFs along $\Delta y$ (*i.e.,* $\Delta x = 0$) for each genotype (in separate columns). Solid curves represent best fit curves. For both rows, blue: cytoplasmic; black: nuclear.

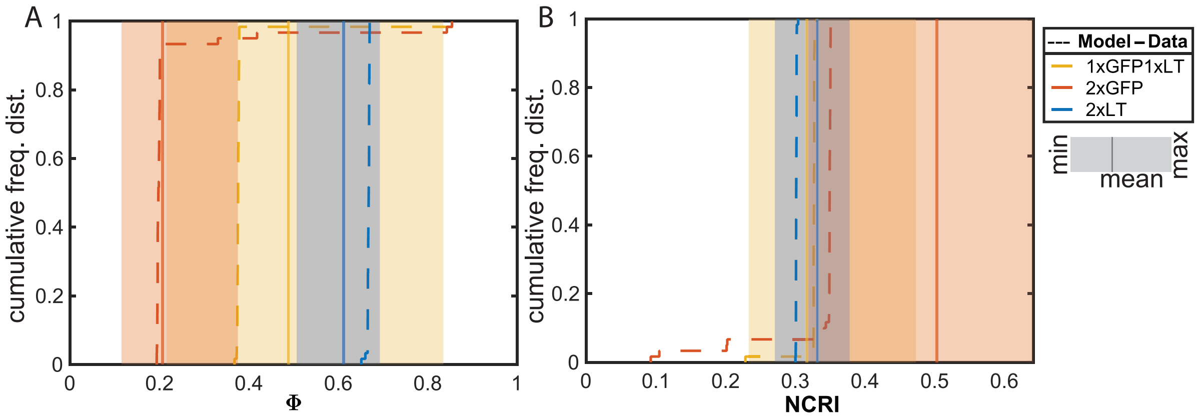

**Figure S4: Model Fits.** Cumulative Frequency distribution of the model predicted $\phi$ (A) and NCRI (B).

**Table S1: IDT gBlock used for CRISPR right homology arm with sequence and description.**

| gBlock Name | gBlock Sequence | Description |
|---|---|---|
| Cact_Right_Homo | GCGCGCTCTTCATATCACTGTAC-<br>CAATTTAAAAATATATTAAGGTGTCCAATGTAAAACTC<br>GATGTTTGTTGCTAATTCTCTCTACCTACGTACGAG-<br>TATATGTGTATGTATTT-<br>GTGTGTGTCTGTTTCTGTATGTTTGTAATTTTTGTATGT<br>GCGTGCGCGTGTGCGTGAATCGAC-<br>CAACTAATCTATTATACAAAATTGAATCATAAACCG-<br>TAGTACTAAGCTAGCTAAATCTAATTGATGTTTTCATG<br>TTCTGACTGAATGAAATAATGTTGCAAAATAAA-<br>TATTTTATTCTACAAAAAAAAATTAG-<br>CAAACAAAAATTAAATGTTGAATTCGATGTTGTAATG<br>GATTGAAAAGAAGGTTTGTAACTCTATTTAGACTT-<br>GAAAGTATGCCCCCCTTTATTGTTTACTTGTTCTTGCT<br>ATGTTGTTTGCTTATTTTATCGCTTAATGAGGCTTTT-<br>GTTCTTGTACCTAACAATTTTGTAA-<br>TATTTTTAATTTAACCCGCTTTTTTCTTGCGAACCTCT<br>TGTTCTTGCAGATGTACGATCGCTTT-<br>GGTGATCCTCGCTATTTTGTCAGCTACAACGGTGG-<br>CAATCCCATGACAGTTGCCGGAGGAGGTGGAGGTGG<br>CATGGCCCAGGTTCAGCTGGTCGAG-<br>TCCGGTGGTGCCCTGGTTCAGCCCGGTGG-<br>CAGCCTGCGCCTGTCCTGCGCCGCCAGCGGTTTCCC<br>AGTGAACCGTTATTCGATGCGCTGGTAC-<br>CGTCAGGCCCCCGGAAAGGAACGCGAG-<br>TGGGTGGCCGGCATGTCCTCCGCCGGCGATCGCTCCA<br>GCTATGAGGATAGCGTCAAGGGCCGTTTCAC-<br>TATCTCCCGCGACGACGCCCGCAACACCGTC-<br>TATTTGCAGATGAACTCCCTGAAGCCCGAGGACACT<br>GCCGTCTACTACTGCAACGTCAACGTCGGATTCGAG-<br>TACTGGGGCCAGGGCACTCAGGTCAC-<br>TGTCTCGTCGTGAATGCCCATCCAATCTGTACATACG<br>ATTTAGACTAAACATAAACATACCTATGAT-<br>TAACACAACTTAAATTCCCGATGTGGTTA-<br>TATCGATGTCCCCAGAAGCACGCCTTTTCTGCCCCAG<br>AATTGTATGCAATATTATTCCTAGACACACAC-<br>GTTCTCAAAATGCATCCAGGAAATGGAAC-<br>CTTATGTACTTACTGTCCCGGCACGTGGGAATCCGAA<br>ACGAATAGAATTCCTTGTGTTAACGCAAAATGTTTA-<br>GAGCGAAATAAAGCATA-<br>CAAGGATGCAGAATGTCAAGCATTTTATTTATACGCA<br>CACAAACTTAACTCTAAGCAA-<br>TAAAAGTAACTGACAAAATTTAATTATTCATA-<br>TATCGTCGGCGGGGGCACATAATCAGAGATTTATCCG | Right homology arm for Cactus CRISPR adding GFP LlamaTag |

| | |
|---|---|
| | TAATTACAAAGTACGAAACATTAAACGA-<br>GAACATAAAAATTGGATATGCATTCAAGAAA-<br>TAATTGTCTAGTCTGGAAAACATACCAGTAGTAAGTA<br>TTGAAGCTTAAACTTAGCAATCAAGGGGG-<br>TAAAAACAGTTCTAAAGGTCTGAGCATATCAAC-<br>GGATGCTCTGTCGGAACACGCTCTGTTTGCCCTTGCT<br>GGTCGAAACTGCCTTCTTGGACGCCAACGGATCGG-<br>GAGCGAAGCAGTTCCAAAGAC-<br>GCAGGGTTTCATCAGCTCCGGCGCTGATCACTGTGCT<br>GCCGTCCGGAGACATGGCCATCTGGA-<br>GAACTCGTGACGTGTGTCCAGTCAAATCGGCTT-<br>GCTTCACCATTGTTGGGTATTTCCAAATGGTCAGTTG<br>GTTGTTAGCAAAACCATGCGCAGAGATCAGCTCCTT-<br>GTAGTGGCGAGAAAAGAGCAGA-<br>GAACAGACCTGCGACTTGGAGTCCACGGATTTCATTA<br>AAGTGCCATTGTTCACATTCCAGAACTT-<br>GATGCAGCGATCGGCGGTGCCGCCTCCAGAGGCTAG<br>AGTACTTGGTTGCCAGGGACACCAGGCCAAGGCAC-<br>GCACTGCAGCTTGATGGTCGTT-<br>GAATTTGGTCAGAAGAGCAAAA |

**Table S2: Primers used for creating CRISPR plasmids with sequences and a brief description.**

| Primer Name | Primer Sequence | Description |
|---|---|---|
| Cact-Target Sense | CTTCGG-<br>CATTCAGGCAACTGTCAT | 5' phosphorylation; forward primer for gRNA for Cas9 target site for insert in Cactus |
| Cact-Target Antisense | AAACATGACAGTT-<br>GCCTGAATGCC | 5' phosphorylation; reverse primer for gRNA for Cas9 target site for insert in Cactus |
| Cact-Intron gRNA Sense | CTTCGTACAGTGCAC-<br>GAAAGTACT | 5' phosphorylation; forward primer for gRNA for Cas9 in-tron site for insertion of DsRed in Cactus |
| Cact-Intron gRNA Antisense | AAACAGTACTTTCGTG-<br>CACTGTAC | 5' phosphorylation; reverse primer for gRNA for Cas9 in-tron site for insertion of DsRed in Cactus |
| Left Homo Arm F | GCGCCACCTG-<br>CAAAATCGCTGGGAA-<br>TAGTATTACAAAGT | Forward primer for CRISPR left homology arm PCR out of genomic DNA |
| Left Homo Arm R | GCGCCACCTG-<br>CAAAATTATCAC-<br>GAAAGTACTTGGGAGAA | Reverse primer for CRISPR left homology arm PCR out of genomic DNA |

**Table S3: Hyperparameters of the Cact-LT/GFP model**

| Hyperparameter | Brief description | Value |
|---|---|---|
| Generations (G) | Number of generations (iterations) the algorithm runs for | 1000 |
| Population ( $\lambda$) | Total number of individuals in each iteration | 150 |
| $\mu$ | | 20 |
| nIslands | Number of islands | 1 |
| migGen | Migration parameter (if nIslands > 1) | -1 |
| $\alpha$ | Smoothing factor | 0.2 |
| $\gamma$ | Recombination parameter | 0.85 |
| pf | Pressure on fitness | 0.45 |
| varphi | Expected rate of convergence | 1 |
| tmax | Maximum time the algorithm will run for | 72h |
| mm | Minimize or maximize | ‘min’ |
| yeslin | Use lin-step | true |
| yesnew | Use newton-step | true |
| nlin | Number of individuals lin-step contributes | 2 |
| nnewt | Number of individual newton-step contributes | 1 |
| nlinPar | | 1 |
| nnewtPar | | 1 |
| startlin | Declare in which generation lin-step should start contributing | 1 |
| Endlin | Declare in which generation lin-step should stop contributing | Last generation |
| Startnew | Declare in which generation newton-step should start contributing | 1 |
| endnewt | Declare in which generation newton-step should stop contributing | Last geneation |
| useFullnewtonstep | | true |
| sortPrevParamters-byError | | false |
| $\beta$lin | | 1 |
| reshufflegen | | 0 |
| restart | | 0 |

Movie Captions:

Movie S1: Timelapse of a whole 1x1x embryo from nc11-nc14, showing the Cact-LT pattern during and between nuclear cycles. The pattern is predominantly cytoplasmic, disappearing when nuclear envelopes dissolve during a nuclear division, and reestablishing after formation of new nuclear envelopes

Movie S2: Timelapse of a control embryo during nc14 at 40x zoom, showing the GFP pattern without the influence of Cact-LT. The pattern is slightly nuclear, in opposition to 1x1x embryos.

Movie S3: Timelapse of a 1x1x embryo from nc11-nc14 in a vertical orientation, detailing the full dynamics around the dorsal-ventral axis. The pattern is the same as described for the whole embryo.

Movie S4: Timelapse of a 1x1x embryo FRAP, covering 10 minutes post bleaching of a single nucleus.

Movie S5: Timelapse of a control embryo FRAP, covering 10 minutes post bleaching of a single nucleus.

Movie S6: Timelapse of a 2x GFP embryo FRAP, covering 10 minutes post bleaching of a single nucleus.

Movie S7: Timelapse of a 1x1x embryo RICS.

Movie S8: Timelapse of a 2x Cact-LT embryo RICS.

Movie S9: Timelapse of a 2x GFP embryo RICS.
